# SuperResNET GUI: model-free single molecule network analysis software achieves molecular resolution of Nup96

**DOI:** 10.1101/2024.03.12.584716

**Authors:** Y. Lydia Li, Ismail M. Khater, Christian Hallgrimson, Ben Cardoen, Timothy H. Wong, Ghassan Hamarneh, Ivan R. Nabi

**Affiliations:** Department of Cellular and Physiological Sciences, Life Sciences Institute, University of British Columbia, Vancouver, BC V6T 1Z3, Canada; Medical Image Analysis Lab, School of Computing Science, Simon Fraser University, Burnaby, BC V5A 1S6, Canada; Department of Electrical and Computer Engineering, Faculty of Engineering and Technology, Birzeit University, Birzeit P627, Palestine; School of Biomedical Engineering, University of British Columbia, Vancouver, BC V6T 1Z3, Canada

**Keywords:** Nup96, nuclear pore, dSTORM, machine learning, network analysis, SuperResNET software

## Abstract

SuperResNET is an integrated machine learning-based analysis software for visualizing and quantifying 3D point cloud data acquired by single molecule localization microscopy (SMLM). The computational modules of SuperResNET include correction for multiple blinking of a single fluorophore, denoising, segmentation (clustering), and feature extraction, which are then used for cluster group identification, modularity analysis, blob retrieval and visualization in 2D and 3D. Using publicly available dSTORM data, we apply a graphical user interface (GUI) version of SuperResNET to nucleoporin Nup96 structures, that present a highly organized octagon structure comprised of eight corners. SuperResNET GUI effectively segments nuclear pores and Nup96 corners based on differential proximity threshold analysis. SuperResNET GUI quantitatively analyzes features from segmented nuclear pore structures, including complete structures with 8-fold symmetry, and from segmented corners. SuperResNET GUI modularity analysis of segmented corners distinguishes two modules at 11.1 nm distance, corresponding to two individual Nup96 molecules. SuperResNET GUI is therefore a model-free tool that can reconstruct network architecture and molecular distribution of subcellular structures without the bias of a specified prior model, attaining molecular resolution from dSTORM data. SuperResNET GUI provides flexibility to report on structural diversity in situ within the cell without model-fitting, providing opportunities for biological discovery.

## INTRODUCTION

Single molecule localization microscopy (SMLM) is a super-resolution microscopy technique that generates 2D or 3D point cloud data by localizing target proteins labeled by isolated fluorophores, providing lateral resolution of 5-20 nm and axial resolution of 10-30 nm using photoactivated localization microscopy (PALM) ^[1, 2]^, stochastic optical reconstruction microscopy (STORM) ^[3, 4]^, and point accumulation for imaging in nanoscale topography (PAINT) ^[5, 6]^. More recently, MINFLUX nanoscopy, which also generates point cloud data by localizing single fluorophores labeling the protein of interest, achieved 1-3 nm 3D resolution ^[7, 8]^. SMLM has been applied to study biological structures including actin, microtubules, nuclear pore complex, clathrin coated pits, caveolae and focal adhesions ^[9, 10]^, providing visualization and reconstruction at the nanometer scale.

While software packages for reconstruction and visualization of SMLM data are well-developed,^[11]^ approaches for quantitative analysis of 2D or 3D point cloud data remain limited. Clustering analysis methods for SMLM include statistical, Bayesian, density-based, correlation-based, tessellation-based, image-based, and machine-learning based approaches ^[12, 13]^. Previously, we applied batch SuperResNET network analysis to SMLM data for caveolin-1 (CAV1) and identified caveolae and three distinct classes of non-caveolar scaffolds ^[14-16]^. Modularity analysis shows that small CAV1 scaffolds combine to form larger scaffolds and caveolae ^[15]^, and convex hull analysis detected changes in caveolae structure due to point mutation in the caveolin scaffolding domain ^[17]^. Here, we present a user-friendly graphical user interface (GUI) version of SuperResNET for application of 3D SMLM network analysis method ^[15, 18]^, providing fast 3D cluster identification and classification from SMLM point cloud data sets.

SuperResNET GUI is robust for denoising SMLM data, segmenting clusters, extracting features of clusters, and using machine learning to classify clusters. Additionally, SuperResNET GUI enables visualization of segmented clusters and retrieves clusters based on selected feature(s).

The nuclear pore complex is used extensively as a reference structure for super-resolution microscopy because of its well-defined structures and abundance in the cell ^[19]^. Approximately 30 different nucleoporins are found in the nuclear pore complex, and electron microscopy has identified the structures of most of the nucleoporins (Supplemental Figure 1) ^[20]^. Here, we evaluate the capability of SuperResNET GUI to analyze publicly available 2D dSTORM data for nucleoporin Nup96 ^[19]^ and demonstrate its ability to effectively segmenting nuclear pores and Nup96 corners. While most nuclear pores are incomplete, SuperResNET GUI is able to extract the size, shape and network features of all segmented blobs and selected complete ones. Modularity analysis of segmented corners distinguishes 2 modules at 11.1 nm distance corresponding to the 2 individual Nup96 molecules, known to be located at 12 nm distance within the nucleopore corners ^[20]^. SuperResNET GUI network analysis of dSTORM data of Nup96 ^[19]^, with a predicted resolution of 13.3±0.1 nm, therefore obtains molecular resolution at 12 nm. Uniquely and in contrast to recent model-based methods, SuperResNET’s analysis modules do not require the user to specify what structure to reconstruct in advance, maximizing the potential for discovery.

## MATERIALS AND METHODS

### Data source

List of localizations of publicly available 2D SMLM data for Nup96 was obtained from Thevathasan et al. ^[19]^. To summarize, U20S cells expressing Nup96-SNAP labeled with Alexa Fluor 647-benzylguanine were fixed and imaged using dSTORM. Localizations generated by custom software written in MATLAB were corrected for drift, merged to remove localizations persistent over consecutive frames, and filtered by localization precision and fitting quality.^[19]^

### SuperResNET GUI

The SuperResNET GUI was built in MATLAB and based on previous SMLM Network analysis software ^[18]^. Here, SuperResNET was used to analyze the publicly available 2D dSTORM Nup96 data referenced above. Proximity thresholds for the analysis were determined using Ripley’s H-function. To remove background and random localization, we constructed a random localization network and extract node degree of each localization at proximity threshold 12 nm, compared the distribution of random network/graph with experimental data and retained localizations with degree greater than the average network degree of the random graph multiplied by a scalar alpha (*a*).

The mean-shift algorithm was used to segment nuclear pores and Nup96 corners in the denoised localizations using kernel bandwidth determined from Ripley’s H-function, and 30 features of each blob (Supplemental Figure 3) were extracted to describe the size, shape/anisotropy, topology, and network features and corresponding statistical features. The 30 features were then used to characterize blob groups using K-means clustering, where 2 groups were identified in segmented nuclear pores and 1 in segmented corners.

Modularity analysis described previously ^[15]^ was applied to segmented corners at proximity threshold 12 nm. Corners with 2 modules were identified and distance between the module centroids was measured.

Additional details on SuperResNET capabilities can be found in the Supplemental Methods section. SuperResNET GUI is available as Supplementary Software. Updated versions of the software can also be found at *https://www.medicalimageanalysis.com/software/superresnet*. The batch analysis version of SuperResNET can be accessed at *https://github.com/NanoscopyAI*. SuperResNET software is compiled for Windows and Mac operating systems. The software manual shows the installation process and provides documentation on how to use the software to analyze biological SMLM data. The software comes with a license for academic use only.

## RESULTS

### SuperResNET graphical user interface

SuperResNET is an integrated software to visualize and quantify 3D point cloud data acquired by single molecule localization microscopy (SMLM). The GUI provides users an easy access to loading data, correction of multiple blinks originating from the same fluorophore, filtering of background noise, segmentation of clusters, quantification of cluster features, grouping based on cluster features, network and modularity analysis of individual clusters as well as 2D and 3D visualization under GUI tabs (Figure 1). Key features of SuperResNET GUI include: **Load data**: loads point cloud data and provides histogram visualization of localization and the associated metadata; **Merge & Network analysis**: removes artifacts from multiple blinks and assesses scale of clustering by Ripley’s H-function; **Filter**: denoises data based on comparison with degree distribution of random network; **Segment**: provides Mean-shift segmentation for blob-like structures and DBSCAN to segment structures with other shapes; **Blob features:** extraction and quantification of 30 features comprising size, shape, topology/hollowness, statistical and network features of segmented blobs and reports histograms of the features; **Group**: assigns group to segmented blobs based on selected features through K-means clustering and reports color-coded histograms of grouped features; **Individual blobs**: visualizes individual blobs with network connection at user-defined proximity thresholds; **Blob modules**: extracts and visualizes modules of interacting molecules at user-defined proximity thresholds for individual blobs; **Blob retrieval**: retrieves representative blobs from each group with visualization of localization, network, and boundaries in 2D and 3D.

**Figure 1:**
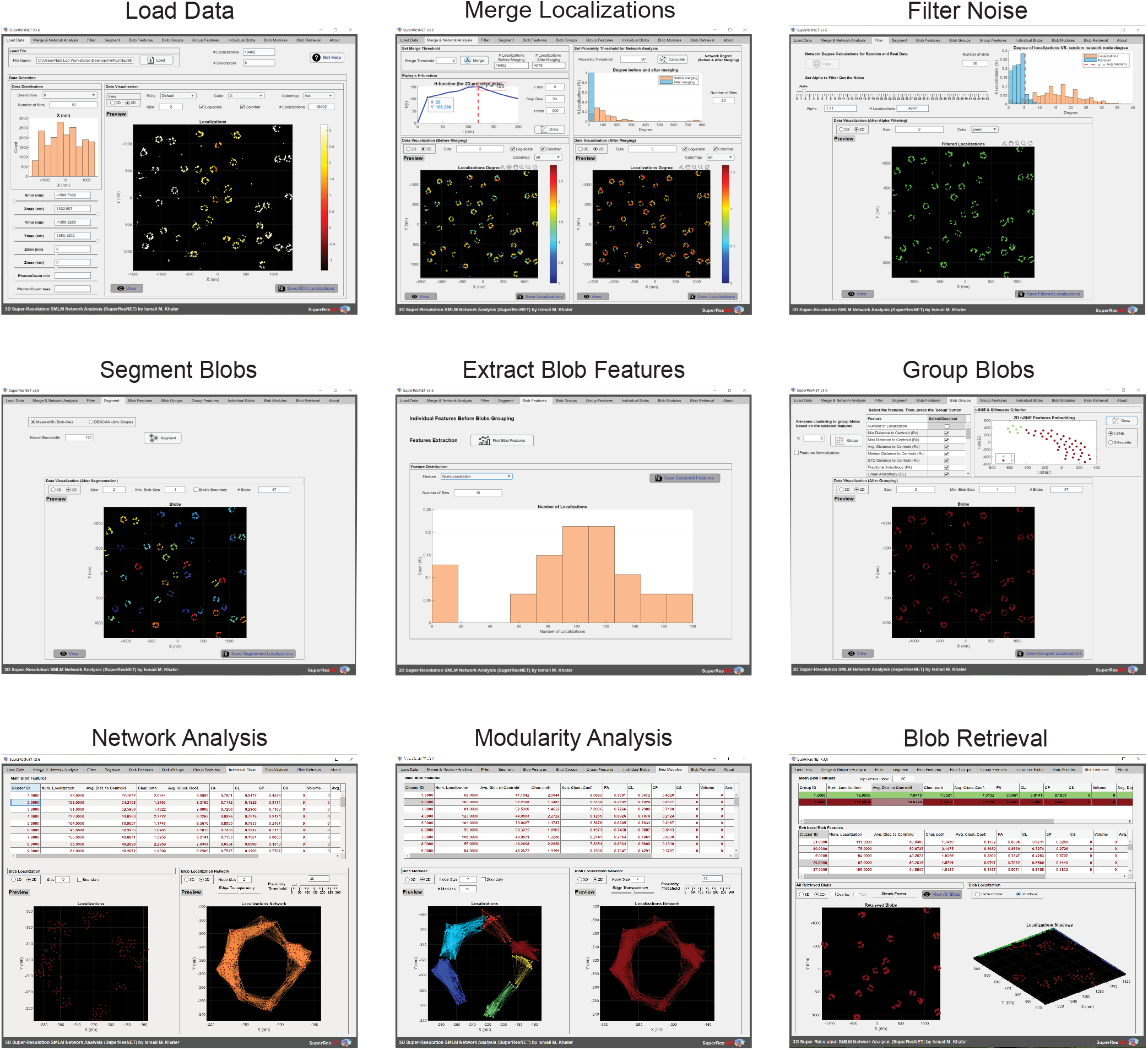
Overview of SuperResNET GUI software. SuperResNET graphical user interface is an integrated software to quantify and visualize 2D/3D point cloud data. The pipeline of SuperResNET includes loading point cloud data, correcting/merging of multiple blinks originating from the same fluorophore, filtering of background noise, segmentation of blobs, extraction of blob features, grouping based on blob features, network and modularity analysis of individual blobs, retrieval of representative blobs as well as 2D and 3D visualization.

### SuperResNET GUI analysis of dSTORM data for Nup96 labeled nuclear pores

We applied SuperResNET GUI to publicly available 2D dSTORM data for Nup96 ^[19]^. The localization event list was loaded into SuperResNET GUI (Figure 2A). Using the SuperResNET GUI merge & network analysis tab, Ripley’s H-function was calculated for the loaded localizations. When plotting Ripley’s H-function *H*(*r*) against radius *r* (Figure 2B), a peak at r=118 nm and an inflection point at r=12 nm were identified. These correspond to the size of nuclear pores (the cluster scale) and Nup96 corners (modules or sub-cluster scale), respectively. In order to best filter the data to retain the corners, we generated a random network and plotted the degree distribution of that localization and that of the same data randomized at proximity threshold 12 nm (Figure 2C). Under the filter tab of SuperResNET GUI (Figure 1), users can define filtering parameter α for denoising, removing localizations with degree lower than *α* × a*verage* degree of random network. For the current dataset, α=13 removed the random localizations (dashed red line in Figure 2C) with 7.8% of total localizations removed and considered not clustered at r=12 nm scale. The effects of filtering are shown in Figure 2D. Localizations outside of nuclear pores were present at α=0 and removed at α=13, whereas the ring of nuclear pore was maintained.

**Figure 2:**
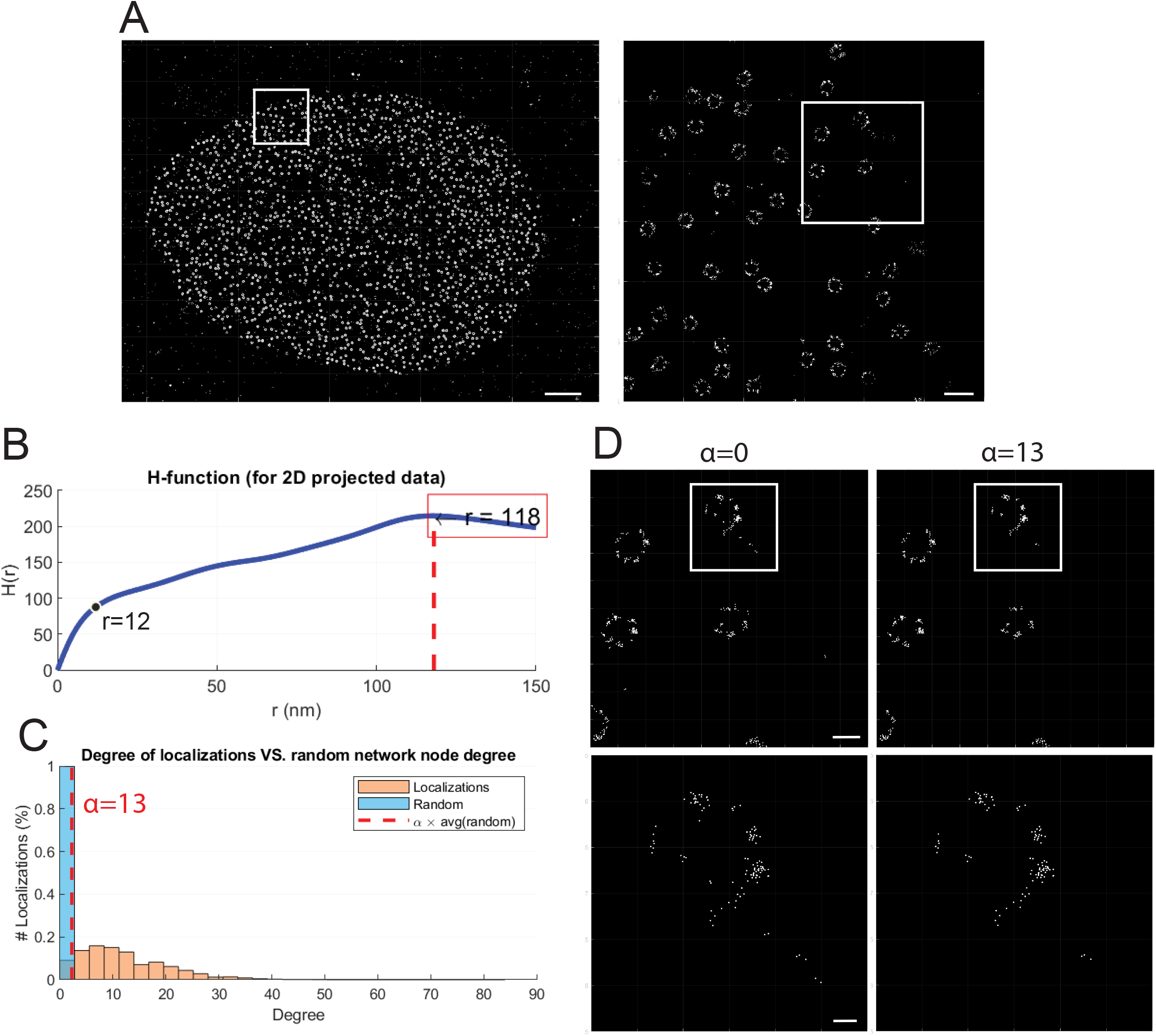
Preprocessing in SuperResNET GUI using merge & network analysis and filter tabs. (A) 2D point cloud data for Nup96 acquired by dSTORM were loaded and visualized in SuperResNET GUI software. Scale bar, 2 μm; zoom, 250 nm. (B) Ripley’s H-function plotted against cluster radii. (C) Degree distribution of random network (blue) and localization (orange). Dashed red line indicates degree value of α × average degree of random network. (D) Effects of filtering at α=0 and α=13. Scale bar, 100 nm; zoom, 25 nm.

Based on the peak of Ripley’s H-function (r=118 nm) (Figure 2B), nuclear pores were segmented using mean-shift algorithm (Figure 3A), and 30 features describing size, topology, hollowness, shape and network were extracted from the segmented blobs. Formal definition of the used features is provided in the Supplemental methods. Feature analysis of the segmented blobs showed that for area, average distance to centroid, characteristic path, modularity, network density, number of localizations, x range and y range, there were two clear peaks (Supplemental Figure 3). We therefore identified 2 groups by K-means clustering using the extracted features (Figure 3B), with 16.6% of blobs classified as Group 1 and 83.4% as Group 2; quantification of features after grouping is shown in Figure 3C. Group 1 blobs contain an average of 10±0.4 localizations per blob, while Group 2 blobs contain 106.8±1.0 localizations per blob. Group 1 blobs are smaller in size compared to Group 2, with X range of 19.9±1.0 nm and 130.3±0.9 nm, respectively. Group 1 blobs also show lower modularity and higher network density than Group 2, while Group 2 blobs are hollower than Group 1 (Figure 3C). This suggests that Group 1 blobs correspond to isolated clusters of Nup96 and Group 2 blobs correspond to more complete nuclear pores.

**Figure 3:**
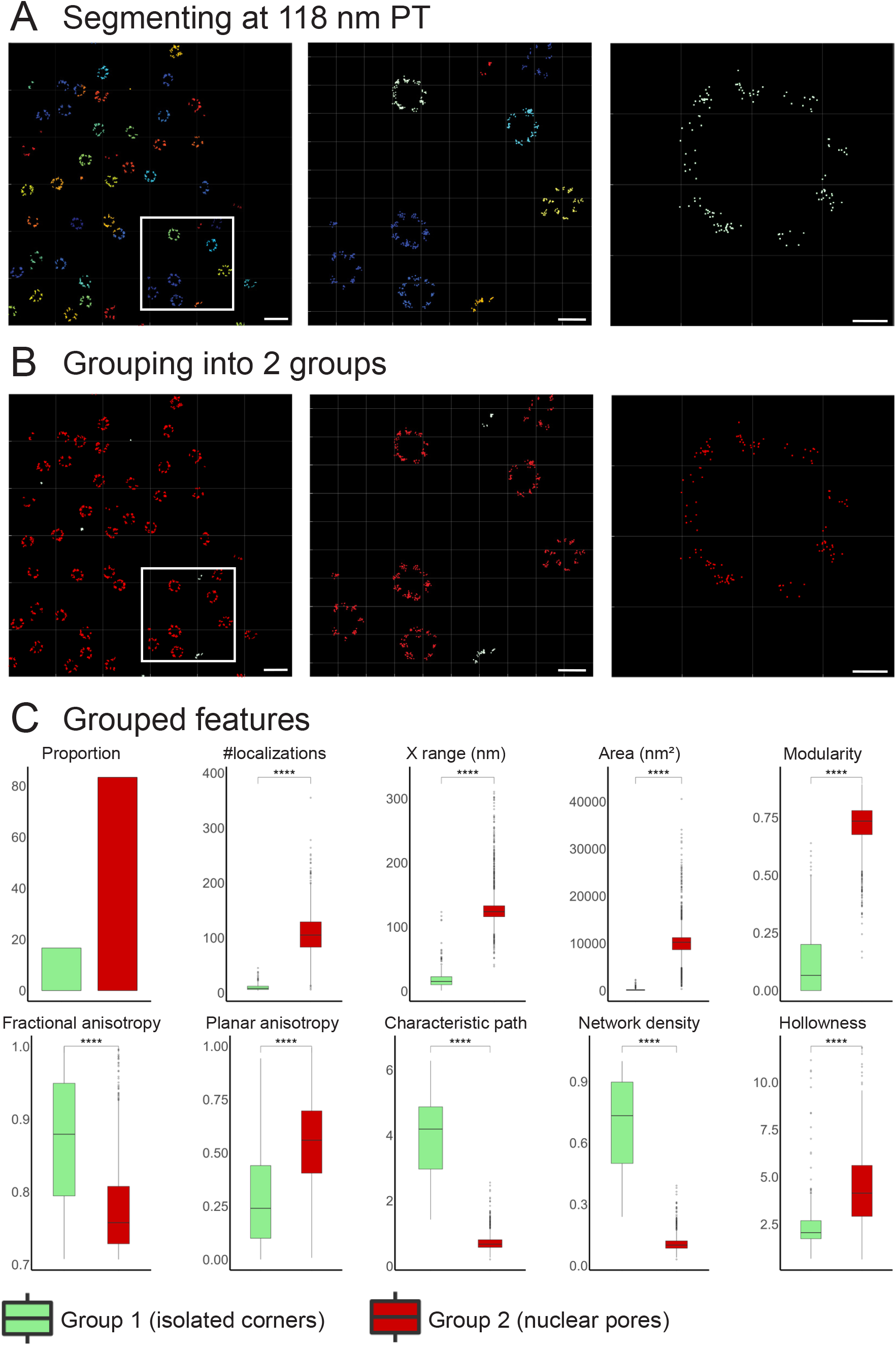
Segmentation and grouping of nuclear pores using segment and group tabs. (A) Segmentation of nuclear pores using mean-shift algorithm. (B) Grouping of segmented blobs using blob features by K-means clustering. Nuclear pores (Group 2, red) and isolated corners (Group 1, green) were classified into two separate groups. Scale bars, 250, 100 and 20 nm (left to right). (C) Features of grouped blobs. Quantification of blob size, anisotropy (shape) and size features (n=293 for Group 1 and n=1469 for Group 2; two-sided t-test, ^****^ *p* < 0.0001). The central line denotes the median value, the box contains the 25^th^ and 75^th^ percentiles, the whiskers marks the 5^th^ and 95^th^ percentiles and circles denote outliers.

The top 49 representative Group 2 blobs with features closest to the mean blob features were retrieved (Figure 4A) using the SuperResNET GUI blob retrieval tab (Figure 1). While most nuclear pores were incomplete and did not show 8 corners, some complete pores can be identified and network connection of two complete nuclear pores at different proximity thresholds (Figure 4B) were visualized using the individual blobs tab (Figure 1). Localizations within the same corners were connected into networks at proximity thresholds 10 nm, adjacent corners were connected at proximity threshold 50 nm, and corners on opposite sides of the nuclear pore were connected at proximity threshold 100 nm. These SuperResNET values correspond to the 42 nm distance between adjacent corners and 107 nm distance between opposing corners of the nucleopore complex, as determined by cryoelectron microscopy ^[20]^.

**Figure 4:**
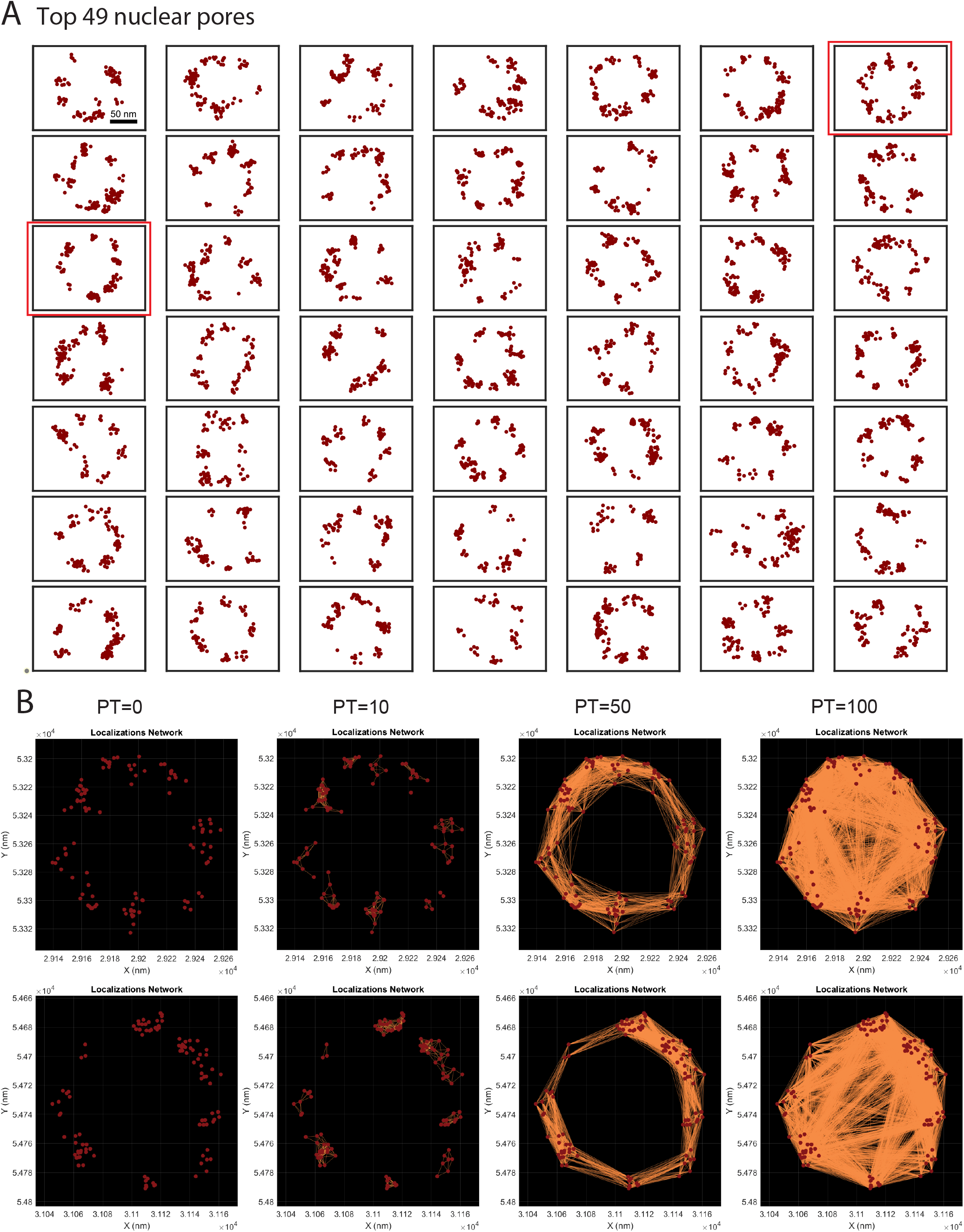
Network connection of representative nuclear pores retrieved using blob retrieval and network analysis tab. (A) Localizations of top 49 representative nuclear pores. Boxed nuclear pores were shown in (B) network connections at proximity threshold 0, 10, 50, and 100 nm.

### SuperResNET GUI identified 2 Nup96 molecules in Nup96 corners by modularity analysis

Based on the inflection point in Ripley’s H-function (Figure 2B), Nup96 corners were then segmented at proximity threshold 12 nm using mean-shift algorithm (Figure 5A). Corners contain 12 localizations on average and are 15.3±0.1 nm in X range. They are low in modularity (0.06) and high in network density (0.77) (Figure 5B), which resemble the features of Group 1 blobs when segmenting at proximity threshold 118 nm. Euclidean distance was calculated using all features to compare segmented corners at proximity threshold 12 nm and previously identified Group 1 and Group 2 when segmented at proximity threshold 118 nm (Figure 5C). The Euclidean distance between segmented corners and Group 1 are highly similar and far from Group 2. This suggests that Group 1 blobs are isolated corners dissociated from complete nucleopore complexes.

**Figure 5:**
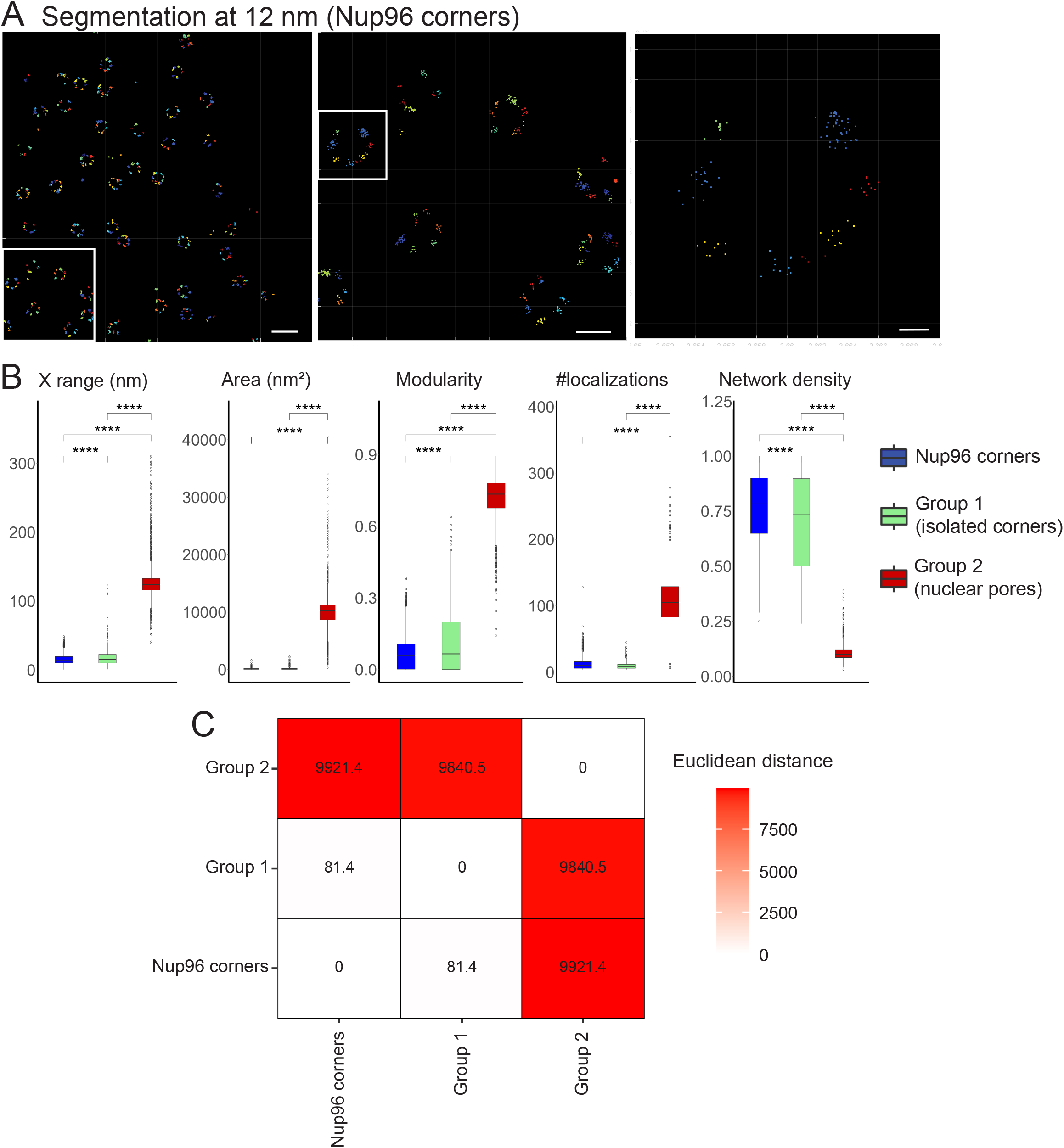
Segmentation of Nup96 corners using segment tab. (A) Segmentation of nucleoporin corners using the mean-shift algorithm. Scale bars, 250, 100 and 20 nm (left to right). (B) Features of segmented corners (blue), Group 1 (green) and Group 2 (red) in segmented pores (n=12636 for corner, n=293 for Group 1 and n=1469 for Group 2; ANOVA with Tukey post-test, ^****^ *p* < 0.0001). The central line denotes the median value, the box contains the 25^th^ and 75^th^ percentiles, the whiskers marks the 5^th^ and 95^th^ percentiles and circles denote outliers. (C) Euclidean distance of the group features of segmented corners and groups in segmented pores.

We then used the SuperResNET GUI modularity analysis tab (Figure 1) to apply modularity analysis to the segmented corners, in which subnetworks of high-density localizations are identified within segmented blobs ^[15]^. Blobs with 2 modules were identified at proximity threshold 12 nm and the distance between module centroids was measured. Those blobs containing fewer localizations show highly variable distance between module centroids (Figure 6A), likely a consequence of poor identification of modules based on few localizations. To identify a threshold for blobs showing low variance of distance between modules, we calculated the index of dispersion of distance between module centroids, plotted the value against number of localizations, and identified the inflection point at 11 localizations (Figure 6B). Blobs with greater than 11 localizations have a mean distance between module centroids of 11.13 nm (Figure 6C), corresponding to the 12 nm distance between Nup96 molecules measured by cryoelectron microscopy ^[20]^. Representative blobs with two modules were visualized using the blob modules tab (Figure 1) and shown in Figure 6D.

**Figure 6:**
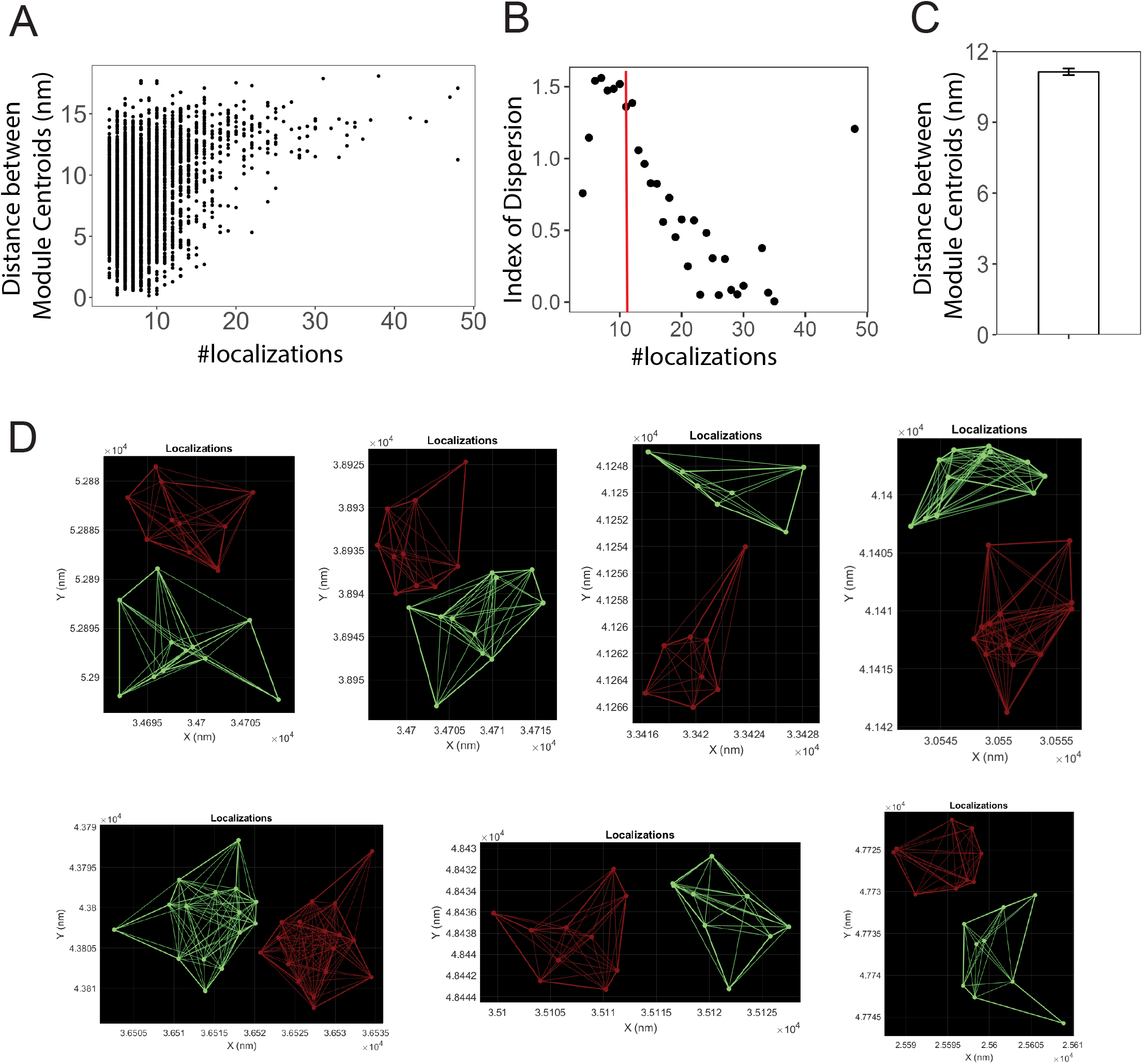
Modularity analysis of segmented Nup96 corners using blob modules tab. (A) Relationship between number of localizations and distance between centroid of 2 modules in Nup96 corners. Blobs with 2 modules were identified at proximity threshold 12 nm. (B) Relationship between number of localizations and variance of distance between module centroids divided by mean. Red line indicates 11 localizations per Nup96 corner. (C) Mean distance between module centroids in blobs with greater than 11 localizations. Error bars denote standard error. (D) Representative blobs with 2 modules.

## DISCUSSION

SuperResNET GUI is a new GUI version of the previously described SMLM network analysis algorithm SuperResNET^[15, 18]^. The algorithm represents a user-friendly approach to filter, segment, group and visualize the structures found within the point cloud of localizations generated by various SMLM approaches, including dSTORM. The GUI incorporates each element of the analysis pipeline into sequential tabs and provides easy access for users to adjust parameters. Here, we show that application of SuperResNET GUI network analysis to publicly available dSTORM Nup96 data is able to reconstruct the 8-fold symmetry of the nuclear pore as well as segment the nucleoporin corners and detect the individual Nup96 molecules at 12 nm distance within the nucleopore corners, essentially attaining molecular resolution from dSTORM data. This analysis highlights the analytic improvements that network analysis brings to interpretation of SMLM data.

SuperResNET GUI provides the ability to segment SMLM datasets at different proximity thresholds, essentially analyzing the extent to which localizations interact at varying distances. SuperResNET GUI includes a Ripley’s H function analysis tool that defines optimal clustering distances. For Nup96, SuperResNET GUI provides the ability to segment and analyze complete nuclear pore complexes as well as the nucleoporin corners, based on the 118 and 12 nm proximity thresholds obtained from Ripley’s H-function analysis. Euclidean comparison of feature of the two groups obtained at 118 nm and the corners obtained at 12 nm shows that the smaller Group 1 obtained at 118 nm proximity threshold closely matches the corners obtained at 12 nm proximity threshold. This suggests that Group 1 blobs represented isolated corners. These isolated corners are far less abundant than complete nuclear pores and are not seen as clusters larger than one corner. Whether they represent corners isolated from nuclear pores fragmentation due to fixation and processing of the samples or isolated corners within the nuclear membrane is not clear. Biological structures, from proteins to DNA to more complex protein complexes are modular in nature. SuperResNET GUI provides the ability to compare the features of multiple structures within the cell and thereby define the relationship between various oligomeric structures derived from interaction between the same component molecule, in this case Nup96.

Similarly, prior SuperResNET analysis of CAV1 defined caveolae and smaller scaffold structures and was able to show structural correspondence between the modules that comprise these different structures ^[15]^. This approach was incorporated into SuperResNET GUI modularity analysis, in which sub-modules of larger structures are identified through increased proximity of subsets of localizations. Here, application of SuperResNET modularity analysis to the Nup96 corners was able segregate clusters of localizations that were ~12 nm apart corresponding to the two individual Nup96 proteins known to be found within nucleoporin corners. The ability to obtain molecular resolution at 12 nm from dSTORM highlights the potential of SuperResNET GUI to define molecular architecture of cellular structures in situ within the cell. Indeed, application of SuperResNET GUI to MINFLUX data sets of clathrin coated pits in the accompanying paper provides highly resolved clathrin pit and vesicle structures ^[21]^.

Nevertheless, determination of molecular localizations from SMLM data remains challenging. A priori knowledge that nucleoporin corners comprise two Nup96 molecules ^[20]^ led to a focus on two modules. Indeed, use of biological knowledge to inform interpretation of SMLM data is critical to guiding interpretation ^[22]^. Prior application of modularity analysis to caveolin-1 clusters was able to report on the number of modules in each structure based on local proximity of localizations ^[15]^. In that case, detected modules reported interaction between presumed CAV1 molecules as application of the SuperResNET merge module reduced local networks, formed due to the well-characterized multiple blinking issue of

SMLM ^[23]^. Multiple blinking occurs due to the fact that the same fluorophore can blink more than once; due to drift between blinking events the multiple blinks do not necessarily coincide, forming a local cluster. Reduction of these local clusters is possible in SuperResNET using the variable threshold merge module. The merge threshold was previously set to 19-20 nm based on prior biological knowledge that approximately 150 CAV1’s formed a caveolae, and encompassed both localization precision of the detected blinks as well as error introduced by primary and secondary F(ab’)^2^ antibody labeling ^[14]^.

As the SNAP-tagged Nup96 data analyzed in the current study was already preprocessed using a temporal method by merging localizations that persist over consecutive frames ^[19]^, we did not merge localizations for this dataset. The fact that some individual Nup96 molecules within corners remained associated with multiple blinks within the localization precision of the acquisition of 13.6 nm ^[19]^ indicates that preprocessing had not effectively removed multiple blinks. Further, as can be seen from the views of the most representative nucleopore complexes (Figure 4A), only few contain eight corners the majority and the majority are incomplete. The combination of multiple blinking and incomplete coverage for this SMLM dataset of a highly defined structure, the nuclear pore complex, is an indication of the challenges faced when reporting on molecular structure with SMLM ^[19]^.

SuperResNET’s spatial approach to multiple blinking is best supported by saturation of labeling and blink acquisition. As can be seen in this study, in which localization merging in SuperResNET GUI was not applied, these retained local clusters due to multiple blinking provide information that enabled modularity analysis of the corners and localization of individual Nup96 molecules. Importantly, a limited numbers of localizations for some corners resulted in poor identification of modules and highly variable reporting of distances between the two modules. Exclusion of corners with few localizations enabled accurate determination of the distance between the two clusters derived from the two Nup96 molecules within a nucleopore corner, a distance that was below the localization precision of the acquisition. This highlights the power of cluster analysis for SMLM data and also the way the multiple blinking “problem” of SMLM can be leveraged to inform on underlying molecular structure.

One approach to correct for incomplete labeling is particle averaging, where multiple identical copies of the structure of interest are combined to reconstruct a “super-particle” with high labeling density and signal-to-noise ratio. However, common template-based methods ^[24, 25]^ are prone to template bias, and methods derived from single particle analysis for cryoelectron microscopy ^[26, 27]^ are not compatible with 3D SMLM data. Recent development of template-free 3D particle fusion methods ^[28, 29]^ overcomes these limitations, but assumes a single consistent structure and is therefore unable to report on structural variation within the dataset. In contrast, with extended acquisition of blinks and more coverage of the imaged structures, SuperResNET provides the user with flexibility to report on structural diversity in situ within the cell without model-fitting, providing opportunities for biological discovery.

