## Supplemental Methods and Figures for "SuperResNET GUI: model-free single molecule network analysis software achieves molecular resolution of Nup96"

### Supplemental material

**Supplemental Methods:** Provides extended detail on the SuperResNET GUI software.

**Supplemental Figure 1:** Schematic of nuclear pore. (A) Electron microscopy density of the nuclear pore complex with the C termini of Nup96 shown in red. (B) Side view and (C) top view of the nuclear pore. Reproduced with permission from Thevathasan et al. [1].

**Supplemental Figure 2:** Histograms of features of segmented nuclear pores (proximity threshold 118) showing two peaks retrieved by blob features tab.

**Supplemental Figure 3:** All features of segmented Nup96 corners (blue), Group 1 (green) and Group 2 (red) in segmented pores (n=12636 for corner, n=293 for Group 1 and n=1469 for Group 2; ANOVA with Tukey post-test, ** $p < 0.01$, *** $p < 0.001$, **** $p < 0.0001$).

**Supplemental Figure 4:** Per-localization feature extraction for noise filtering, retaining clustered localizations, and segment the clustered localizations. (A) Simulated data that consists of 5 bivariate Gaussian clusters and random noise. (B) The network construction at PT defined by Ripley's H-function. (C) The node degree features. (D) The generated random data that correspond to the simulated data. (E) The random graph constructed using the same PT used in the simulated data graph construction. (F) The degree features of the random graph. (G) The filtered localization shows the clustered nodes after getting rid of the noise. (H) The degree distribution for both the simulated and random graphs and the alpha user-control to remove the noisy localizations shown on (G). (I) Ripley's H-function for the simulated data in (A) shows the scale of clustering.

---

1. J. V. Thevathasan, M. Kahnwald, K. Cieslinski, P. Hoess, S. K. Peneti, M. Reitberger, D. Heid, K. C. Kasuba, S. J. Hoerner, Y. Li, Y. L. Wu, M. Mund, U. Matti, P. M. Pereira, R. Henriques, B. Nijmeijer, M. Kueblbeck, V. J. Sabinina, J. Ellenberg, J. Ries. *Nat Methods.* 2019, 16, 1045.

### SuperResNET GUI Software

**Super**-**res**olution **Net**work analysis (**SuperResNET**) is a software package that contains multiple computational modules for analyzing 3D Super-resolution single molecule localization microscopy (SMLM) data. SuperResNET is a powerful tool for 3D SMLM data analysis, pre- and post-processing, quantification, and visualization.

**Note 1**. SuperResNET can be used to process 2D data by padding the Z dimension with zeros matching the X and Y columns. SuperResNET can be used to process/visualize the 2D data. However, some of the features will be calculated to zeros in Blob Features module (e.g., volume) as these features are designed for 3D data only.

**Note 2**. SuperResNET can be installed on a personal computer. It can be used to process the whole field of view (FOV) or a cropped region of interest (ROI). However, for big SMLM data with large FOV (e.g. > 54 microns) or a very large number of localizations (e.g. > 1 million), then we recommend using a machine with large memory (e.g. 32GB, 64GB, or even larger). SuperResNET can be used to crop ROIs to improve runtimes.

**Note 3**. SuperResNET is very fast. However, for segmenting big-data using the mean-shift [2, 3] algorithm, it might take minutes to hours depending on the size of the data. SuperResNET also supports using DBSCAN [4] algorithm to segment biological structures (i.e. clusters) of various shapes.

---

2. K. Fukunaga, L. Hostetler. *IEEE Transactions on information theory.* 1975, 21, 32.
3. D. Comaniciu, P. Meer. *IEEE Transactions on Pattern Analysis & Machine Intelligence.* 2002, 1, 603.
4. M. Ester, H. P. Kriegel, J. Sander, X. Xu. *InKdd.* 1996, 2, 226.

### Methods

SuperResNET processes SMLM data by extracting features at various levels. Per-localization network features extracted for every localization for denoising, retaining the clustered localizations, and cluster segmentation tasks. On the other hand, per-blob/cluster features extracted for every cluster for cluster characterization and class identification (e.g., using unsupervised machine learning) tasks. In the below sections, we are describing how SuperResNET calculates the various types of features and uses them for several computational tasks (e.g., filtering, segmentation, class identification).

### Per-localizations Features Extraction & Tasks

**Network/Graph Construction**

Given SMLM point cloud data $P = \{p_1, p_2, \dots, p_L\}$, where $L$ is the total number of acquired localizations ($i.e., |P| = L$). Each $p_i$ is a spatial location for the reconstructed event/fluorophore/molecule (i.e., fluorophore localization for the raw data, or molecular localization for the corrected events via the merge algorithm) such that, $p_i = (x_i, y_i, z_i)$ for 3D coordinates at the nano-meter scale. An undirected graph/network $G = (V, E)$ is constructed based on a proximity threshold ($PT$). Where $V$ is the graph vertices/nodes and $E$ is the set of graph edges. The graph nodes represent the localizations (i.e., $|V| = L$). The graph edges represent the interaction between every two pair of nodes/localizations in the graph $G$. The number of edges in the graph is related to the parameter $PT$ and the distribution of the points $P$ in the space of the cell. The user can set $PT$ to construct various graphs for the same point set $P$. $PT$ can be set based on the scale of the clustering. For example, an edge $e_{ij} \in E$ is created between point $p_i$ and point $p_j$ if $|p_i - p_j| \leq PT$. Notice that the interaction of node $p_i$ and node $p_j$ is reciprocal (i.e., the distance from node $p_i$ to $p_j$ is the same as the distance from node $p_j$ to $p_i$). Hence, we have undirected graph/network. More formally, $e_{ij} = \begin{cases} 1, & |p_i - p_j| \leq PT \\ 0, & otherwise \end{cases}$

We here describe a method of setting $PT$ for blob-like structures based on Ripley's H-function. However, the user can manually set $PT$ to construct a graph for other shape structured data (i.e., $PT$ can be set as the width of the tubular structures). As a rule of thumb, we can construct a network to extract the homogenous clusters in the data by setting $PT$ such that ($\frac{r}{2} \leq PT \leq r$). For heterogeneous clusters, $PT = \frac{r}{2}$. Where $r$ is the cluster scale that maximized the Ripley's H-function $H(r)$.

Supplemental Figure 4 shows the process of constructing a network for simulated data (Supp. Fig. 4A). The simulated data consists of five bivariate Gaussian clusters. Each cluster generated with $\sigma = 5\ nm$, covariance matrix $COV = \begin{bmatrix} \sigma^2 & 0 \\ 0 & \sigma^2 \end{bmatrix}$, and number of localizations = 25. The means $\mu_1\ to\ \mu_1$ for each cluster are, $\mu_1 = [0, 0], \mu_2 = [40, 40], \mu_3 = [-40, -50], \mu_4 = [0, 60],\ and\ \mu_5 = [60, -40]$. A random noise with uniform distribution is added to the generated clusters to mimic the real SMLM data (Supp. Fig. 4A). To construct a network for the simulated data, Ripley's H-function is used to find the $PT$ (Supp. Fig. 4J). The constructed network is shown in Supplemental Figure 4B.

#### Degree Feature Extraction

For undirected graphs such as our network, the degree is a node feature that is defined as the number of incident edges to that node. Formally, the degree of a node $i$, $deg_i = \sum_{j \in V} e_{ij}$. Supplemental Figure 4C shows the degree of every node for network constructed in Supplemental Figure 4B.

#### Noise Filtering

In the context of SMLM data, the noisy localizations can be defined as the background and non-clustered localizations that appear in the data due to imaging artifact and monomeric molecules. Noise filtering is defined as labeling every localization using a binary label $l$, such that $l = \{S, N\}$. Where node $p_i$ gets the label $l_i = S$ if it is signal/clustered node and gets $l_i = N$ if it is a noise.

We designed a filtering method that is based on the network features of the real SMLM data and the corresponding random graph features to model the noisy localizations and label them accordingly. The features of the clustered localizations should be different than the noisy ones and get a different label. To do so, we constructed a random graph with the same number of nodes for the simulated data (Supplemental Figure 4E). The random data should be distributed and generated to occupy the same space of the simulated/real data (Supplemental Figure 4D). A random graph should be constructed with the same $PT$ (i.e., defined according to Ripley's H-function) used in the real/simulated data (Supplemental Figure 4E). The unweighted degree measure is calculated for every localization of the random graph (Supplemental Figure 4F). In the real data, we retain the localizations with the features that different than the random graph (i.e., getting label $l = \{S\}$ for clustered nodes). To give the user a control over the filtering process, we used a scalar $\alpha$. The node $p_i$ is labeled as $l_i = S$ if $deg_i > \alpha \times \overline{degR}$. Where, $\overline{degR} = \frac{\sum_{j=1}^{L} degR_j}{L}$, $\overline{degR}$ is average node degree of the random graph, and $degR_j$ is the per-node degree feature of the random graph. Supplemental Figure 4H shows how $\alpha$ is set to retain the clustered localizations and filter out the noisy localizations (Supplemental Figure 4G).

#### Segmenting Clustered Localizations

After the filtering step, the blobs/clusters are obtained via the segmentation process. Cluster segmentation task is the process of decomposing the signal labeled localizations $P_S$ (i.e., $\forall\ P\ with\ l = \{S\}$) into smaller disjoint (non-overlapping) subsets, such that their union covers the whole set $P_S$. Each subset (a cluster/blob $b_i$) represents an underlying macromolecular biological structure. Formally, let us define $C$ as the clustering operation applied on the set $P_S$. $C$ takes every localization $p_i \in P_S$ and label it with a cluster ID such that every localization $p_i$ has one membership to one and only one cluster ID. Usually, the number of clusters (biological structures) is not known and some algorithms (e.g., mean-shift) have no prior assumption about the maximum number of clusters in the data. Hence, the assignment process starts assigning the neighboring localizations (based on the kernel bandwidth) to cluster ID =1. A new cluster ID is generated when there is no more localizations can be added to all the previous IDs. The cluster IDs keep incrementing until all the localizations get assigned to cluster IDs.

Assume after convergence we have K blobs/cluster IDs denoted $B$ ($b_j, j = 1,2, \dots, K$), we write the mapping $C: P_S \rightarrow B\ as\ C(p_i) = b_j \in B,\ \cup_j b_j = P_S, b_i \cap b_k = \emptyset, \forall j, k$, where ∪ and ∩ are the union and intersection operations respectively. Supplemental Figure 3I shows the five clusters segmented using mean-shift algorithm with kernel bandwidth defined using the Ripley's H-function (Supplemental Figure 4J). As a rule of thumb, we can select the kernel bandwidth to be less than or equal to the $r$ value that maximized Ripley's H-function.

### Per-Blob/-Cluster Features Extraction

After the clustered localizations get segmented into blobs/clusters, a set of features are extracted for every individual blob. The features comprise geometrical shape, topology, hollowness, statistical, network, and size for every blob. Given a blob $b_i \in B$, $b_i = \{p_1, p_2, \dots, p_N\}$, where all the localization of $b_i$ have the same cluster ID. The following features can be used to describe the blob. For more details about the definitions of features and their formulae, we refer the reader to [5, 6] for the network features, and to [7] for the shape feature (FA, CL, CS, and CP).

1. The number of localizations feature is the total number of localizations belonging to every segmented blob $b_i$ such that all the localizations have the same cluster ID. **The number of localizations** is $|b_i| = N$.
2. Distance to centroid feature is used to quantify how hollow the blob is by finding the distance of every localization to the center of the blob. Let $c_i$ represent the center of the blob $b_i$, $c_i = \frac{\sum_{j=1}^{N} p_j}{N}$. The distance to centroid of localization $p_j \in b_i$ is $d_j = |p_j - c_i|$. The vector that contains the distances of all localizations of blob $b_i$ to its centroid is $Rc = [d_1, d_2, \dots d_N]$. We find the following features, **average distances to centroid** is $\overline{Rc_i} = \frac{\sum_{j=1}^{N} d_j}{N}$.
3. The **minimum distance to centroid** is $MinRc_i = \min\ (Rc)$.
4. The **maximum distance to centroid** is $MaxRc_i = \max\ (Rc)$.
5. The **median distance to centroid** is $MedRc_i = \text{median}\ (Rc)$. The median for localizations distances to centroid of the vector $Rc$ is the *middle* number when the distances are ordered from smallest to greatest.
6. The **standard deviation of the distances to centroid** is $StdRc_i = \sqrt{\frac{\sum_{j=1}^{N}(d_j - \overline{RC_i})^2}{N}}$.
7. The ellipsoidal shape features for every 3D blob are extracted based on the eigenvalue decomposition of the covariance matrix of the localizations of blob $b_i$. The eigenvalues ($\lambda_1, \lambda_2, \lambda_3$) of the blob $b_i$ are extracted using principal component analysis (PCA). The **fractional anisotropy (FA)** feature is used to describe the degree of anisotropy of the blob diffusion. $FA_i =$

5. Newman, M. E. J. *SIAM review.* 2023, 45, 167.
6. M. Rubinov, O. Sporns. *Neuroimage.* 2010, 52, 1059.
7. C. F. Westin. *Proc. ISMRM'97.* 1997.

$\sqrt{\frac{3((\lambda_1-\bar{\lambda})^2+(\lambda_2-\bar{\lambda})^2+(\lambda_3-\bar{\lambda})^2)}{2(\lambda_1^2+\lambda_2^2+\lambda_3^2)}}$, where the eigenvalue mean $\bar{\lambda} = \frac{(\lambda_1+\lambda_2+\lambda_3)}{3}$. FA values between [0, 1]. A zero value means the blob is isotropic and a value of one means that the blob is elongated along one of the directions and fully restricted along the other two directions.

8. The **linear anisotropy (CL)** feature is used to describe when the elongation of the blob is only along one direction (i.e., $\lambda_1 >> \lambda_2 \cong \lambda_3$) (e.g., cigar shape). $CL_i = \frac{\lambda_1-\lambda_2}{\lambda_1}$.
9. The **planar anisotropy (CP)** feature is used to describe when the blob shape is restricted to the plane defined by two equal eigenvalues (i.e., $\lambda_1 \cong \lambda_2 >> \lambda_3$) (e.g., pancake shape). $CP_i = \frac{\lambda_2-\lambda_3}{\lambda_1}$.
10. The **spherical anisotropy (CS)** feature is used to describe when the blob diffusion is isotropic (i.e., $\lambda_1 \cong \lambda_2 \cong \lambda_3$). $CS_i = \frac{\lambda_3}{\lambda_1}$.
11. The **volume** feature is calculated using the convex hull of the Delaunay triangulation for the 3D blob.
12. A network/graph $G_i = (V_i, E_i)$ is constructed for the blob $b_i$. Where $V_i$ is the set of nodes for the blob $b_i$, $|V_i| = N$, and $E_i$ is the set of edges connecting the nodes when using the $PT$ for the construction of graph $G$. That is, an edge $e_{uv} \in E_i$ is created between point $p_u \in V_i$ and point $p_v \in V_i$ if $|p_u - p_v| \leq PT$. More formally, $e_{uv} = \begin{cases} 1, & |p_u - p_v| \leq PT \\ 0, & otherwise \end{cases}$. The graph $G_i$ is used to extract a set of network features. The node degree feature is extracted for every localization belonging to the blob $b_i$. The degree of a node $p_u$, $deg_u = \sum_{v \in V_i} e_{uv}$. The vector that contains the degree of all localizations of blob $b_i$ is $Deg = [deg_1, deg_2, \ldots deg_N]$. The **average degree** feature for blob $b_i$ is $\overline{deg_i} = \frac{\sum_{j=1}^{N} deg_j}{N}$.
13. The **maximum degree** feature for blob $b_i$ is $maxDeg_i = \max(Deg)$.
14. The **minimum degree** feature for blob $b_i$ is $minDeg_i = \min(Deg)$.
15. The **characteristic path** feature is defined as the average shortest path length between all pairs of nodes in the network. $charPath_i = \frac{1}{N}\sum_{u \in V_i} L_u$, where $L_u$ is the average distance between node $u$ and all nodes in blob $b_i$.
16. The node eccentricity feature ($ecc_u$) is defined as the maximum distance between node $u$ and all nodes in blob $b_i$. The vector that contains the eccentricity of all localizations of blob $b_i$ is $Ecc = [ecc_1, ecc_2, \ldots ecc_N]$. The **average network eccentricity** feature for blob $b_i$ is $\overline{ecc_i} = \frac{1}{N}\sum_{u \in V_i} ecc_u$.
17. The minimum network eccentricity is **network radius** feature. The network radius of the blob $b_i$ is $radius_i = \min(Ecc)$.
18. The maximum network eccentricity is **network diameter** feature. The network diameter of the blob $b_i$ is $diameter_i = \max(Ecc)$.
19. The clustering coefficient feature of an individual node $p_u \in V_i$ (local clustering coefficient) is defined as the fraction of triangles around $p_u$ (the fraction of $p_u$'s neighbors that are also neighbors of each other). The clustering coefficient of node $p_u$, $cc_u = \frac{2\, tri_u}{deg_u(deg_u-1)}$, where $tri_u$ is the number of triangles around node $p_u$, $tri_u = \frac{1}{2}\sum_{v,h \in V_i} e_{uv}e_{uh}e_{vh}$. The vector that

contains the clustering coefficient of all localizations of blob $b_i$ is $CC = [cc_1, cc_2, \dots cc_N]$. The **average clustering coefficient** feature for blob $b_i$ is $\overline{cc}_i = \frac{1}{N}\sum_{u\in V_i} ecc_u$.

20. The **maximum clustering coefficient** feature of the blob $b_i$ is $maxCC_i = \max(CC)$.
21. The **minimum clustering coefficient** feature of the blob $b_i$ is $minCC_i = \min(CC)$.
22. The **network density** feature of the blob $b_i$ is the fraction of actual connections/edges to potential connections. The network density of the blob $b_i$ is $netDen_i = \frac{E_i}{PC_i}$, where $PC_i$ is the potential connections of the undirected network of the blob $b_i$, $PC_i = \frac{N(N-1)}{2}$.
23. The **network transitivity** feature is a variant of clustering coefficient that measures the ratio of triangles to triplets in the whole network. The transitivity of the blob $b_i$ is $trans_i = \frac{\sum_{u\in V_i} 2\, tri_u}{\sum_{u\in V_i} deg_u(deg_u - 1)}$.
24. The **network modularity** feature is a measure of the strength of subdividing the network into nonoverlapping modules/communities. The modularity for the network of the blob $b_i$ is $mod_i = \sum_{u\in M}[e_{uu} - (\sum_{v\in M} e_{uv})^2]$, where the network for the blob $b_i$ is fully subdivided into a set of nonoverlapping modules $M$, and $e_{uv}$ is the portion of all the edges that connects nodes in module $u$ with nodes in module $v$.
25. The **optimized network modularity** feature is the optimal modular structure for a given network that is estimated based on an optimization algorithms to speed up the modularity computations. The optimization algorithms is trying to find the nonoverlapping modules that maximize the number of intra-module edges, and minimize the number of inter-modules edges.
26. The **x range** feature is a measure for the spread of the localizations along the x-dimension of the blobs space. The x range of the blob $b_i$ is $x\_range_i = max(x\_corrdinates_i) - \min(x\_corrdinates_i)$.
27. The **y range** feature is a measure for the spread of the localizations along the y-dimension of the blobs space. The y range of the blob $b_i$ is $y\_range_i = max(y\_corrdinates_i) - \min(y\_corrdinates_i)$.
28. The **z range** feature is a measure for the spread of the localizations along the z-dimension of the blobs space. The z range of the blob $b_i$ is $z\_range_i = max(z\_corrdinates_i) - \min(z\_corrdinates_i)$.
29. The **hollowness** feature is a measure on how spread out the localizations are from the centroid relative to their mean distance from it. The hollowness of blob $b_i$ is $hollowness_i = \frac{mean(Rc)}{std(Rc)}$.
30. The **blob area** feature is the 2D (x, y) area of a blob.

### SuperResNET Software Features Summary

1. SuperResNET supports loading SMLM point cloud data from various microscopes (e.g., dSTORM) with different file formats (e.g., non-text binary, text and ascii formats).
2. SuperResNET enables the user to quickly visualize the data distributions as well as select the required localizations according to some of the associated meta-information (e.g., photon count).

    1. Support saving high-quality plots. Also, it supports interacting with the plots (e.g. put the cursor on a point to get its coordinates and other info).
    2. The software is enabled with preview plots for fast analysis.
3. SuperResNET is equipped with four methods to crop an ROI.
    1. Default ROI method by dragging sliders to select X, Y, and Z values for a specific region.
    2. Rectangular ROI where the user can interactively draw a rectangular region on top of the cell to select an ROI.
    3. Elliptical ROI where the user can interactively draw an elliptical region on top of the cell to select an ROI.
    4. Polygonal ROI where the user can use free-hand to interactively draw any shape region to select an ROI.
4. SuperResNET can be used to interactively visualize the selected SMLM data before and after processing in 2D/3D with several colormaps for the localizations or any of the associated meta-information descriptors. The user can zoom-in/out, rotate, save the plots, etc. quickly.
5. SuperResNET can be used to correct for multiple blinking of a single fluorophore imaging artifact.
6. SuperResNET supports using Ripley's H-function to detect the scale of clustering globally when dealing with spherical/circular-like shape clusters.
    1. Ripley's H-function gives the user a heuristic about how to select the optimal parameters for constructing the network proximity threshold (PT) and segmenting the blobs when using mean-shift (kernel bandwidth).
7. SuperResNET can be used to construct a proximity thresholded undirected network/graph and extract unweighted degree measure as a per-localization feature for further analysis of the localizations.
    1. Ripley's H-function can be used to return the global scale for the clustering (r). As a rule of thumb, we can construct a network to extract the homogenous clusters in the data by using a PT as ($r/2 \leq PT \leq r$). For heterogeneous clusters, $PT = r/2$.
8. SuperResNET can denoise the SMLM data by using the per-localization degree feature to retain the clustered localizations and get rid of the background and non-clustered ones.
9. SuperResNET supports two segmentation methods by assigning a unique cluster ID for every localization. Each localization can be assigned to one and only one cluster ID.
    1. Mean-shift algorithm to segment blob-like clusters (aka blobs).
    2. DBSCAN to segment any other shape structures.
10. SuperResNET can be used to extract per-cluster/blob features/descriptors. The features comprise shape, size, volume, hollowness, topology, and network measures of every individual blob. Also, the user can visualize the features distributions for the blobs.
11. SuperResNET can be used to group the blobs into K classes based on their features. The embedded features can be visualized using t-SNE. The user can select the features used for grouping and whether to normalize these features.
12. SuperResNET supports visualizing the distributions of the individual features after grouping the blobs into K classes. Also, SuperResNET supports the pairwise features visualization.

13. SuperResNET supports browsing the blobs' features and their corresponding localizations.
    1. After segmenting and identifying the blobs' groups, SuperResNET visualizes the blob's features in a tabular way with the consistent color of the identified blobs.
    2. The user can sort, filter, and navigate the blobs features.
    3. The user can interactively show the blob's localizations, the 2D/3D boundary at various shrink factors.
    4. The user can show the blob's network and can interactively change the network's PT of one blob.
    5. SuperResNET visualizes the blob with colors consistent with the blob's groups.
14. SuperResNET supports modularity analysis for the blobs. The user can interactively select a blob and visualize its modules/communities of interacting molecules at various PTs. The user can visualize the modules' 2D/3D boundaries at various shrinking factors.
15. SuperResNET is enabled with a blob retrieval module that can be used to retrieve TOP similar blobs from each group according to their similarities to their average group's features.
    1. The user can visualize and overlay the retrieved blobs in various ways (e.g., overlay the localizations, the 2D/3D boundaries).
    2. The user can apply principal component analysis (PCA) to visualize the retrieved blobs consistently.
    3. The user can visualize the individual blobs and their 2D projections (shadows).
16. SuperResNET software can be shared with other researchers and for various biological applications.
17. SuperResNET supports saving the raw and processed data in CSV format. It also allows saving the blobs, their features, labels, etc. This is important if the user decided to use third-party software (e.g. statistical analysis) to further analyze of the data.
18. SuperResNET supports command-line use with SuperResNET-Batch, and is capable of loading the data output produced by SuperResNET-Batch for visualization.

### SuperResNET Installation

SuperResNET is a cross-platform software that can be run on Microsoft Windows and MacOS. It requires the MATLAB Runtime installer. We packaged the SuperResNET application such that the MATLAB Runtime downloaded from the web. i.e. generates an installer that downloads the MATLAB Runtime and installs it along with the SuperResNET. We explain the installation process for Microsoft Windows and Mac machines in the following documentation.

https://www.medicalimageanalysis.com/software/superresnet

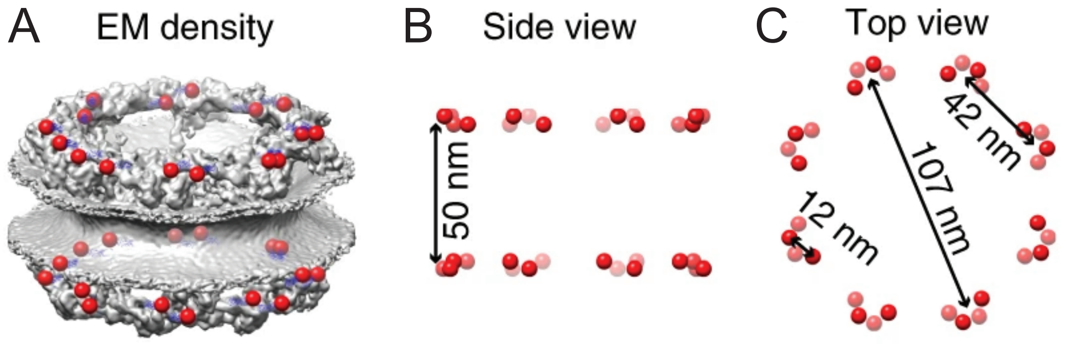


Supplemental Figure 1

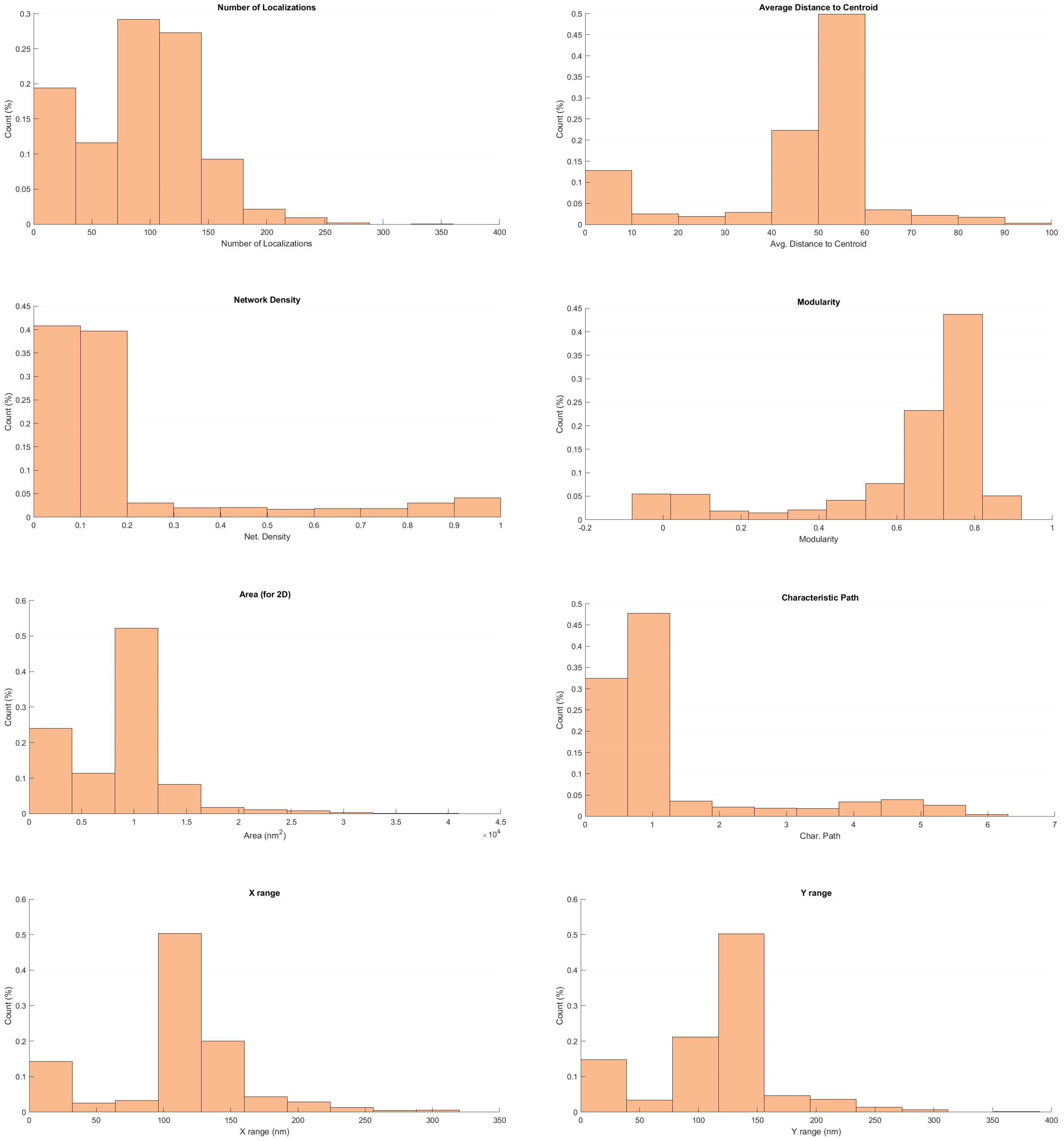


Supplemental Figure 2

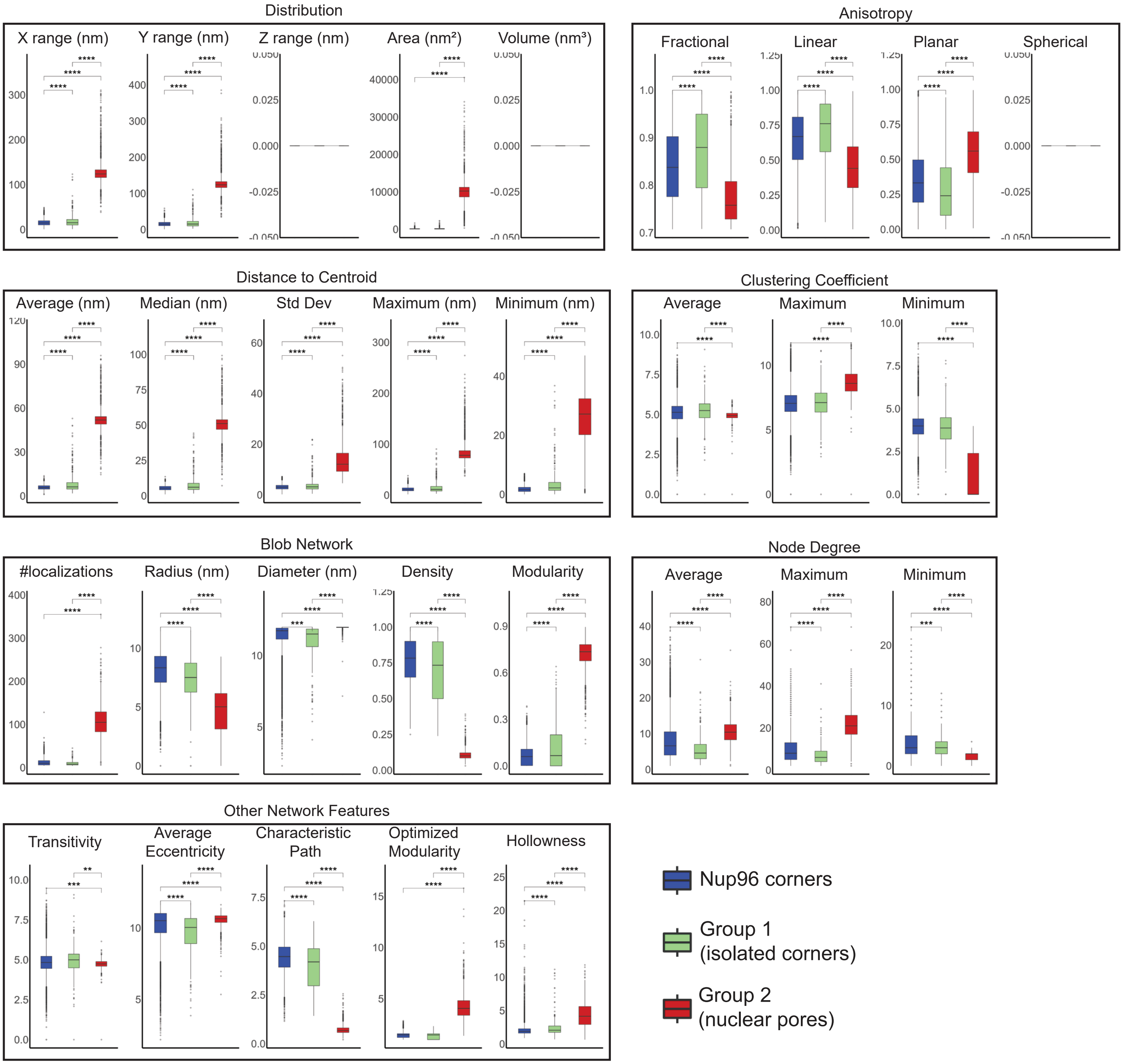


Supplemental Figure 3

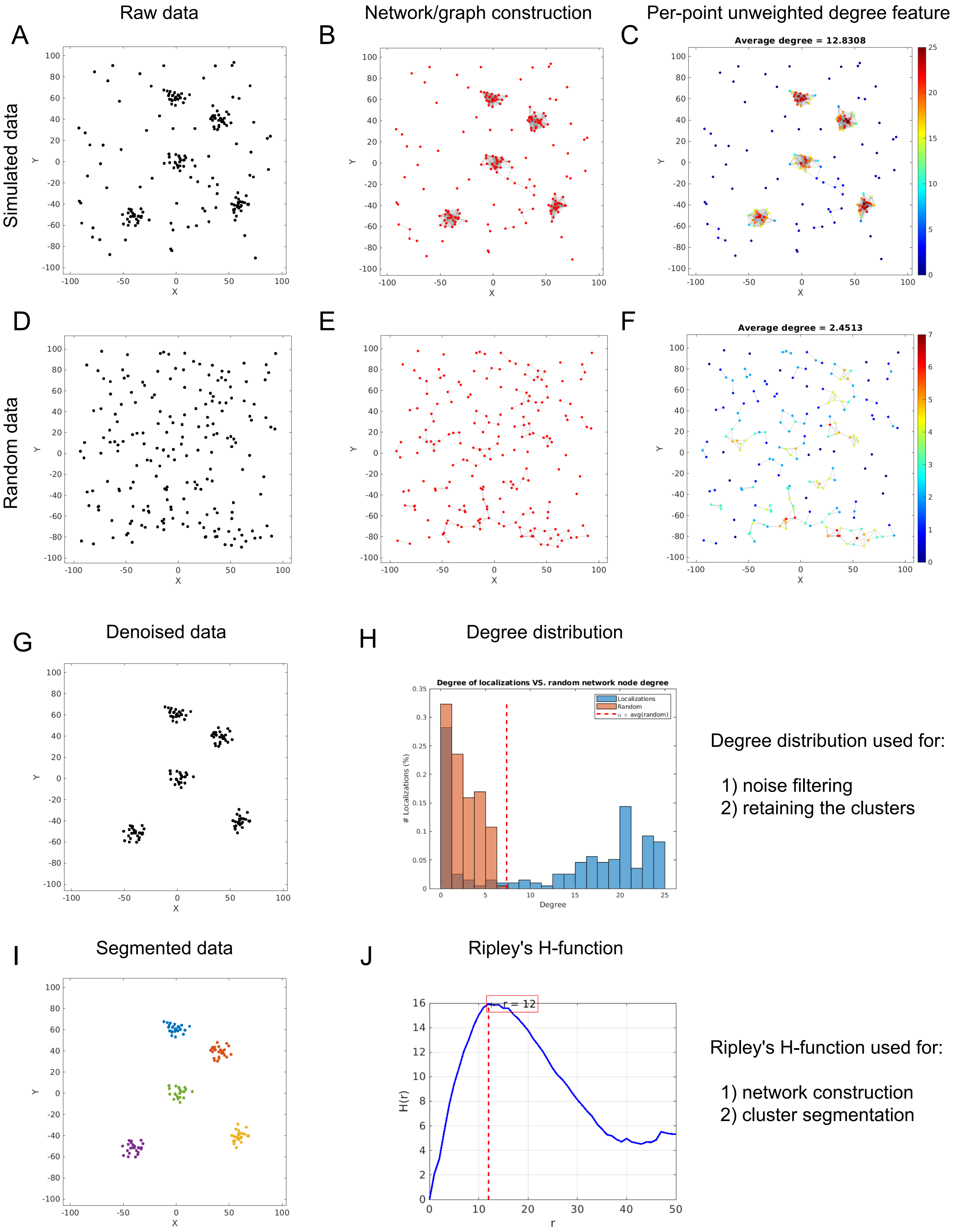


Supplemental Figure 4
